# A codon model for associating phenotypic traits with altered selective patterns of sequence evolution

**DOI:** 10.1101/2020.03.04.974584

**Authors:** Keren Halabi, Eli Levy Karin, Laurent Guéguen, Itay Mayrose

## Abstract

Changes in complex phenotypes, such as pathogenicity levels, trophic lifestyle, and habitat shifts are brought on by multiple genomic changes: sub- and neofunctionalization, loss of function, and levels of gene expression. Thus, detecting the signature of selection in coding sequences and associating it with shifts in phenotypic state can unveil the genes underlying complex traits. Phylogenetic branch-site codon models are routinely applied to detect changes in selective patterns along specific branches of the phylogeny. These methods rely on a pre-specified partition of the phylogeny to branch categories, thus treating the course of trait evolution as fully resolved and assuming that transitions in phenotypic states have occurred only at speciation events. Here we present TraitRELAX, a new phylogenetic model that alleviates these strong assumptions by explicitly accounting for the uncertainty in the evolution of both trait and coding sequences. This joint statistical framework enables the detection of changes in selection intensity upon repeated trait transitions. We evaluated the performance of TraitRELAX using simulations and then applied it to two case studies. Using TraitRELAX, we found an intensification of selection in the SEMG2 gene in polygynandrous species of primates compared to species of other mating forms, as well as changes in the intensity of purifying selection operating on sixteen bacterial genes upon transitioning from free-living to an endosymbiotic lifestyle.

## Introduction

The operation of selection on heritable traits leaves distinct signatures in the genes that code for them. These include, for example, depletions in amino acid changing mutations in genes whose function is crucial (Drory et al. 2004; Gertow et al. 2004; Melamed et al. 2004) or rapid accumulation of mutations in pathogenic genes evading the host immune system of drugs (Endo et al. 1996; Burch and Chao 1999). Therefore, analyzing selection fingerprints at the molecular level while considering phenotypic changes can reveal changes in the selective patterns upon repeated phenotype transitions and the identity of the genes that are associated with the phenotype. The ongoing advances in high-throughput sequencing and increasing efforts to collect phenotypic trait data (e.g., Kattge et al. 2011; Ashman et al. 2014; Parr et al. 2014; Rice et al. 2015), provide the opportunity to detect associations between evolutionary patterns at the genomic level and whole-organism phenotypic traits. Specifically, detecting associations between such traits and selective forces operating at the codon level can provide insight on the locations of functional domains in coding regions that shape the traits of interest and reveal functionalities of unknown genes. With the increased availability of large-scale genome sequence data, the demand for comparative methods for detection of such functionalities is growing. However, such methods are scarce (Nagy et al. 2020).

The nature of selection acting on a protein-coding gene can be revealed by computing the rate ratio between non-synonymous (amino acid altering) and synonymous substitutions, *ω*. Initial codon models (Goldman and Yang 1994; Muse et al. 1994) incorporated a single *ω* parameter, thus reflecting the assumption of a single selective pressure that operates across the entire sequence, be it purifying (*ω* < 1), neutral (*ω* = 1) or positive (*ω* > 1). Further developments integrated multiple *ω* classes into site-models, thereby allowing variation in the selective regime across codon sites (Yang, Nielsen 2000). Moreover, branch-site models (Yang and Nielsen 2002), in which the selective pressure can vary not only across positions, but also among branches of the phylogeny, can be used to detect site-specific changes of selective patterns across the phylogeny based on a prior partition of the branches into distinct categories (often termed background, *BG*, and foreground, *FG*, corresponding to the ancestral and derived states, respectively).

To date, branch-site models are often used to detect selective signatures at the codon level based on phenotypes of study. For example, using the branch-site model of Yang and Nielsen (2002), the color vision in butterflies and primates have been shown to be associated with positive selection in several sites of the opsin gene (Frentiu et al. 2007), and an evidence of connection between rice domestication and elevated *ω* in several genes has been found (Lu et al. 2006). Furthermore, the mating system in primates has been associated with positive selection in the NYD-SP12 gene that is involved in formation of the acrosome during spermatogenesis (Zhang et al. 2007). In evening primroses, positive selection has been associated with sexual reproduction in the chitinase gene, which takes part in construction of the cell wall and defense against pathogens (Hersch-Green et al. 2012).

The branch-site model RELAX (Wertheim et al. 2015) is designed to detect shifts in selection intensity in a predefined set of *FG* branches relative to *BG* branches. Intensification of selection (either purifying or positive) may be indicative of a shift into more stressful conditions while relaxation of selective pressure may be the result of release of functional constraints upon phenotype transition. The latter may indicate loss of function (Wu et al. 1986; Go et al. 2005) or upcoming lineage extinction (Moran 1996). Analysis using this model allows distinguishing between three scenarios concerning the *FG* branches: (1) intensification of selection, reflected in *ω* values moving further away from 1 in the *FG* branches compared to the *BG* branches; (2) relaxation of selection, with *ω* values shifting towards 1; and (3) no change in selection intensity. To achieve that, RELAX incorporates three *ω* parameters, *ω*_0_ ≤ *ω*_1_ ≤ 1 ≤ *ω*_2_ that represent the site classes of the *BG* branches. The difference in the magnitude of selective pressure between the *BG* and *FG* branches is modeled using a selection intensity parameter *k,* such that each of the three *ω* values of the *BG* branches are raised by the power of *k* to obtain the selective pressures of the *FG* branches. Consequently, *k* < 1 implies relaxation of selection and *k* > 1 implies intensification of selection. Using RELAX, several phenotypes have been shown to be associated with a change in selection intensity at the codon level. In orchids, heterotrophic lifestyle has been associated with relaxed purifying selection on the plastid genome (Feng et al. 2016; Roquet et al. 2016; Wicke et al. 2016), and in rodents there is evidence of intensified selection in subterranean species, compared with fossorial species (Tavares and Seuánez 2018).

A notable shortcoming of existing branch-site models is the requirement for a prior specification of branches of the examined phylogeny into branch categories (e.g., *BG* and *FG*). Based on the observed phenotypes of the extant species, a partition of the branches is usually determined by reconstructing the ancestral phenotypic states using the maximum parsimony or the maximum likelihood principles. Either way, the obtained partition is assumed to represent the phenotypic history of the trait across the phylogeny. However, considering a single partition disregards any uncertainty in the reconstructed evolutionary history of the trait. In addition, such an approach unrealistically forces trait transitions to occur simultaneously with speciation events (i.e., at internal nodes of the tree), while in reality they could occur anywhere along a branch, and could lead to underestimation of the number of trait transitions. Consequently, any misspecification of the branch assignments could potentially result in failure to detect changes in selection patterns, as well as high false positive rate (Lu and Guindon 2014).

To account for uncertainty in trait evolution and for possible associations between the processes of molecular and phenotype evolution, several methods have been developed, in which phenotypic changes are explicitly modelled and analyzed jointly with sequence data. CoEvol (Lartillot and Poujol 2011) is designed for the analysis of continuous phenotypic traits, while TraitRate (Mayrose and Otto 2011), TraitRateProp (Levy Karin et al. 2017) and the method of O’Connor and Mundy (2009) are designed for discrete phenotypes. These methods use a rate matrix for nucleotide or amino acid substitutions to either compose a single phenotype-genotype rate matrix (O’Connor and Mundy 2009) or multiply it by a relative rate parameter according to the history of the phenotype (Mayrose and Otto 2011; Levy Karin et al. 2017). Conceptually, using a single phenotype-genotype rate matrix for the analysis of codon data is a straight-forward extension of O’Connor and Mundy’s model but this approach can result in inconsistent phenotypic reconstructions (as described by Levy Karin et al. 2017). Both TraitRate and TraitRateProp are inadequate for the analysis of codon data due to their assumption that the entire rate matrix is scaled upon trait transition, and thus cannot extract the differential effect on synonymous and nonsynonymous substitutions. Recently, Jones et al. (in press) extended this approach to codon sequences for the detection of adaptive changes in selective regime in association with a phenotype, rather than changes in the selection intensity, which is the focus of the current study.

Here, we present TraitRELAX, which enables the detection of changes in the selection intensity upon transitions between phenotypic states. TraitRELAX considers many possible trajectories of trait evolution (“histories”), weighted by their probabilities (Fig. 1). By doing so, TraitRELAX alleviates the need to rely on prespecified branch partitions and allows trait transitions to occur anywhere along a branch. Using extensive simulations, we measure the sensitivity and specificity of TraitRELAX and examine the accuracy of the inferred parameter estimates under a range of data scenarios. We then demonstrate the use of the method by detecting evidence of relaxation in several house-keeping genes of γ-proteobacteria upon transitioning to endosymbiotic lifestyle, and an intensification of selection in the SEMG2 gene, involved in sperm competition in polygynandrous (i.e., multi-male multi-female mating system) primate lineages.

**Figure 1.**
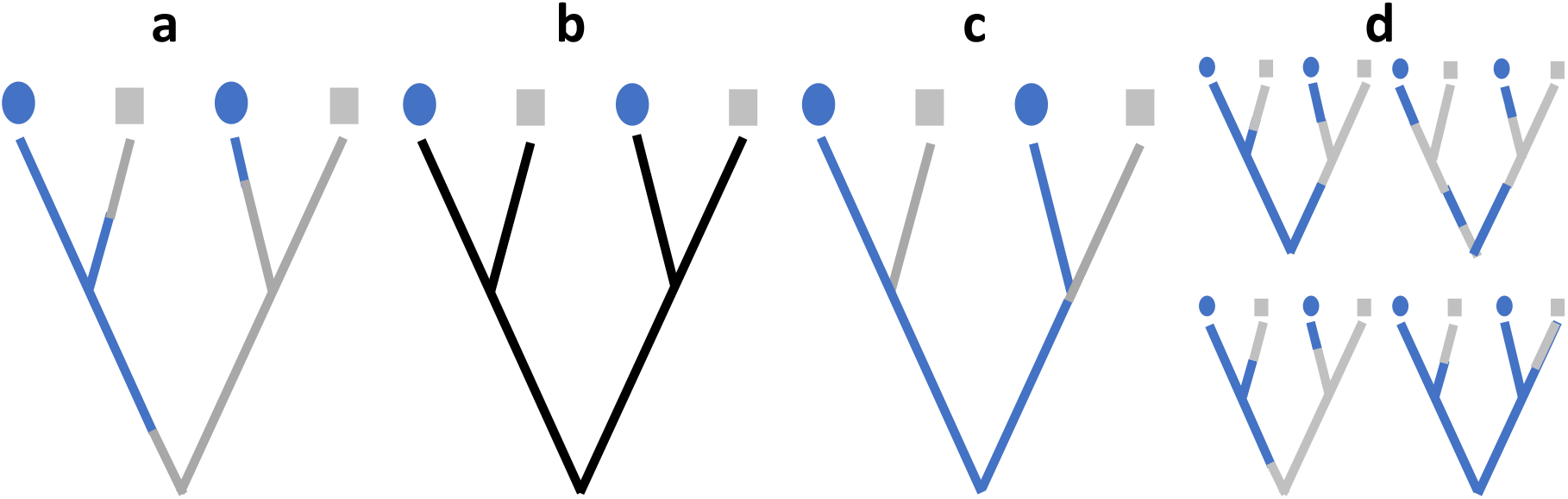
Selection patterns in codon sequence evolution. (a) The true history of a binary trait (square / circle) and the true partition of the branches into categories. (b) Codon models of the whole genome or site families do not allow selective regimes to vary across the phylogeny, assuming a time homogenous selection pattern. (c) Branch-site models allow the selective regime to vary between the branches of the tree, according to *a*-priory transitions pattern. (d) The suggested trait-related codon model co-infers the evolution of the trait, incorporating uncertainty in its history by integrating over multiple possible histories, weighted by their probability.

## Materials & Methods

TraitRELAX is a joint probabilistic model of phenotypic (termed throughout ‘character trait’) and coding-sequence evolution. The input consists of coding sequence data (*D_S_*) in the form of a multiple sequence alignment (MSA), character data (*D_c_*) describing the trait states of the extant species, and a tree with specified branch lengths (T). The model integrates two continuous time Markov processes – one describing the evolution of the character trait (*M_C_*) and the other is a branch-site model that describes the evolution of the coding sequences (*M_S_*). By considering both processes in a joint statistical framework, TraitRELAX can detect relaxation or intensification of selection at the codon level in association with the character evolution. In our branch-site model, branch partition is determined based on the history of the trait of interest, where branch categories are mapped to different character states. Thus, in case of a binary trait, there are two branch categories ‘0’ and ‘1’.

### Character trait evolution

TraitRELAX considers character traits with two possible states coded as either ‘0’ or ‘1’. *M_C_* is defined by the rate matrix *Q_C_*:

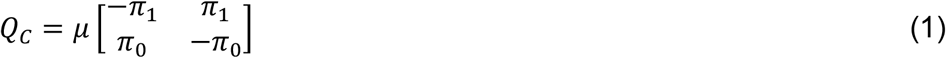

where *π*_1_ = 1 – *π*_0_ represents the rate of change from state ‘0’ to ‘1’ and *μ* is a scaling parameter designated to adapt the branch lengths of *T* to the expected number of character changes. The model described here is not limited to binary character traits but can be applied generally for any number of discrete states using the appropriate rate matrix (Lewis 2001), but in such cases the model would contain additional free parameters.

### Coding sequence evolution

The TraitRELAX sequence model (*M_s_*) is based on the branch-site model RELAX (Wertheim et al. 2015). This model consists of six rate matrices *Q_bm_*, one for each combination of branch category *b* ∈ {0,1} and site class *m* ∈ {0,1,2}, represented by the parameter *ω_m_*:

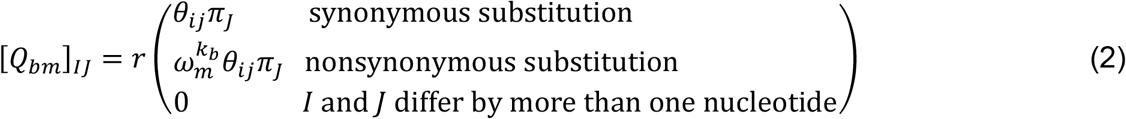

The diagonal elements are determined by the constraint that each row in *Q* sums to zero. Above, *r* represents a scaling parameter that is used to convert the branch lengths of the input tree to units of expected number of codon substitutions; *π_J_* denotes the equilibrium frequency of codon *J*, and *θ_ij_* denotes the rate of substitution from nucleotide *i* to nucleotide *j*, which can be parameterized based on any time-reversible nucleotide substitution model (e.g., Tavaré 1986, Tamura 1992).

Throughout this study we used the K80 nucleotide substitution model (Kimura 1980), which incldues a single parameter describing the transition to transversion rate bias. The *k_b_* parameter represents the selection intensity operating on branch category *b* relative to branch category 0, and thus *k_0_* = 1.

Given a branch partition *B*, the likelihood at site 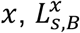, can be computed by averaging over the conditional probabilities 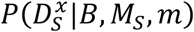 of observing the sequence data at site 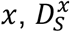, in each site class *m*. For each site class, two rate matrices, *Q*_0*m*_ and *Q*_1*m*_, are alternately used according to the branch assignments in *B*. Considering all site classes, the likelihood thus becomes:

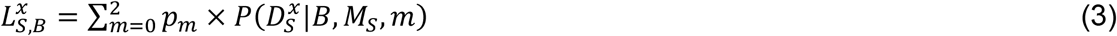

where *p*_0_ and *p*_1_ are free parameters, describing the fractions of sites associated with site classes 0 and 1, respectively, and *p*_2_ = (1 – *p*_0_ – *P*_1_) those with site class 2.

### Joint likelihood computation

The likelihood of the combined model is the joint probability of *D_c_* and *D_s_* given all model parameters and the phylogeny:

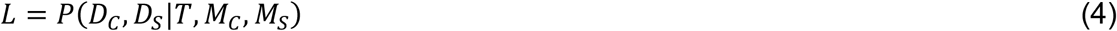

which can be decomposed to:

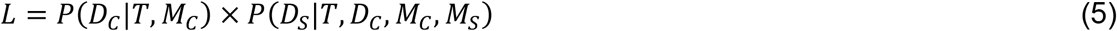

where the first term, denoted as *L_C_*, is the likelihood of *M_C_* and is computed using the pruning algorithm (Felsenstein 1981). The second term, denoted *L_S|C_*, is the likelihood of *M_S_*, conditioned on *M_C_* and *D_C_*. Under these settings the computation of *L_S|C_* requires knowledge of the character state in each part of *T*. This history is generally unknown, but the probability of a given history *P*(*h|T,D_C_,M_C_*) can be computed based on *D_C_* and *M_C_*. Thus, the likelihood of *M_S_* for site 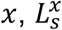 is computed by integrating over all possible character histories, weighted by their probabilities:

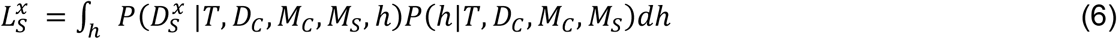

Omitting parameters that do not affect the probability of 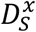 and *h*, we obtain

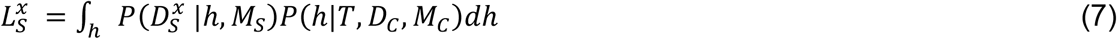

The computation of 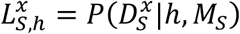 follows from Equation 3. Of note, in the branch partition induced by *h*, character transitions are allowed to occur anywhere along a branch, not necessarily at the internal nodes of the tree. Thus, some branches of the input phylogeny may be divided into several segments, each mapped to a different branch class (in such cases, each point of a trait transition along a branch defines an internal node with a single descendent lineage). Finally, assuming independence of sequence sites, the likelihood of the sequence model becomes:

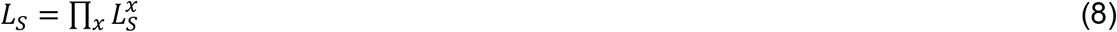

### Approximations

The number of possible character histories is infinite so full integration of Equation 7 is infeasible. Two alternatives were thus used to approximate the full integration in Equation 7: an exhaustive approach and an expected history based approach. In the exhaustive approach, we perform importance sampling using stochastic mappings (Nielsen 2002) and replace the integral by a summation over *N* mappings, each with a probability of 1/*N*:

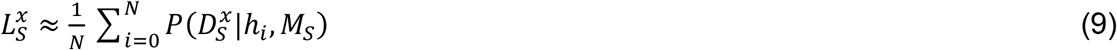

The likelihood computation detailed above requires *N* computations of the sequence likelihood, which is computationally exhaustive. Alternatively, in the expected history based approach, we use a single history *E*(*h*) that represents the expected amount of time spent in each character state along each branch over all possible histories:

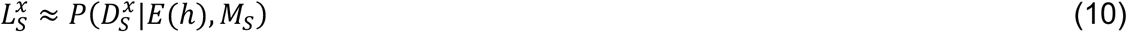

We compute the expected history *E*(*h*) following the rewards method of Minin and Suchard (2008) (see Supp. Text 1). We used the expected history based approach throughout as it proved superior to the exhaustive approach both in terms of runtime as well as robustness (Supp. Text 2).

### Estimating parameters

The TraitRELAX model includes a total of 19 parameters, assuming the K80 nucleotide substitution model is used and that codon frequencies are estimated using the F3×4 model. The nine codon frequencies are estimated from the observed sequence data, while other parameters are estimated using a maximum likelihood optimization procedure that is divided into two stages that are reiterated until convergence: (1) searching for the character model parameters, given a fixed set of sequence model parameters, and (2) searching for the sequence model parameters, given a fixed set of character model parameters. The first stage yields an assignment for the character model parameters, with which the expected character history *E*(*h*) is computed. In this stage, the local search is conducted using either a sequential one-dimensional search (Brent 1974) or a two-dimensional search (Hestenes and Stiefel 1952). In the second stage, the branch partition induced by *E*(*h*) is used for likelihood computations. In this stage, the values for the sequence model parameters are being searched simultaneously using the conjugate gradient method (Hestenes and Stiefel 1952).

### Testing for a trait-related change in selection intensity

A null model, in which the selection intensity parameter, *k*, is fixed at 1 (i.e., imposing the same selection intensity throughout the tree, with no effect of the character trait) can be compared to an alternative model, in which *k* is free to vary (different intensities associated to trait states). Since the models are nested, a likelihood ratio test (LRT) can be used, with the distribution of twice the log-likelihood difference between the two models approaching a 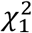 distribution with infinite amount of data under null conditions. In phylogenetic contexts, however, data size is not well defined and is rarely large enough, such that likelihood-ratio test statistics are not always well behaved (Goldman 1993; Whelan and Goldman 1999). As a more exhaustive alternative, the distribution of the likelihood ratio was also estimated using parametric bootstrap. Specifically, sequence data were generated by simulating sequence evolution along the input phylogeny using the parameter estimates of the null model. Given the original character data at the tips of the tree and the simulated sequence data, the maximum likelihood values of the null and alternative models were computed. This procedure was repeated 100 times to produce a distribution for the difference in the maximum log-likelihood values obtained between the two models under null conditions. The critical value for the LR statistic was determined as the 95th percentile of this empirical distribution for a significance threshold of *α* = 0.05. In the simulation study, rather than simulating artificial data sets based on the maximum likelihood estimates obtained by each inference of the null simulated data set, we simulated 200 replicates under the null model with the same parameter values with which the data sets were simulated (except for the selection intensity parameter *k* which was fixed at 1).

### Simulating data sets under the TraitRELAX model

Simulations were used to investigate the sensitivity and precision of our method and to assess its accuracy in inferring the selection intensity parameter *k*. We first generated random trees with different numbers of taxa (16, 32, and 64) according to a birth–death process using INDELible (Fletcher and Yang 2009) with default parameters (speciation rate 0.3 and extinction rate 0.1). We then simulated character evolution along a given tree using Bio++ (Guéguen et al. 2013). As part of these simulations, the exact locations of character state transitions were recorded, thereby yielding branch partition based on the simulated (i.e., true) character histories. This partition was then used to simulate codon sequences based on the sequence model of TraitRELAX using INDELible (Fletcher and Yang 2009). Sequence data were simulated with various numbers of positions (*l* = 150, 300, and 600), representing a typical range of protein lengths (Zhang 2000). Unless otherwise stated, all simulations were conducted with total branches length of 4, the parameters of the character model set to *π*_0_= 0.5 and *μ* = 8 and parameters of the sequence model set to *ω*_0_ = 0.1, *ω*_1_ = 0.8, *ω*_2_ = 2, *p*_0_ = 0.5, *p*_1_ = 0.4 and the transition/transverion rate ratio *kappa* = 2. Different values of the selection intensity parameter were simulated: *k* = 0.2, 0.5, and 0.8 represent relaxation of selection along lineages with character state 1, *k* = 1.2, 1.6, and 2 represent intensification of selection, and *k* = 1 represents no trait-related change in selection intensity. For each simulated scenario, 50 independent replicates were generated (each based on a different tree), with the exception of cases with *k* = 1, for which 200 replicates were generated. The larger number of replicates in the null case was used to determine the empirical threshold for the LR bootstrap-based comparison.

### Biological data analyses

TraitRELAX was used to examine the evolution of Semenogelin II (SEMG2) in 24 primate species with respect to changes in the mating system. Character state assignments (1 = polygynandrous, 0 = other systems) were based on published literature (Dixson 1997; Dixson and Anderson 2002; O’Connor and Mundy 2009). Nine species were assigned as polygynandrous (multi-male:multi-female) and 15 were assigned as ‘other’ (encompassing nine species that are monogamous, single-male:single-female; five species that are polygynous, single-male:multi-female; and one that is polygamous). Sequence data of the SEMG2 gene was collected from previous papers (Ulvsbäck and Lundwall 1997; Jensen-Seaman and Li 2003; Hurle et al. 2007; Roan et al. 2011; Isshiki and Ishida 2019). The accession of the *Otolemur garnettii* was excluded due to exceedingly larger sequence length. In case more than one accession per species was available, a representative accession was selected (the one with maximal pairwise similarity relative to all other accessions). Finally, the assembled sequences were aligned with RevTrans version 2.0b (Wernersson and Pedersen 2003) with default parameters. Positions with more than 90% gaps across sequences were filtered out. The primate tree topology was obtained from TimeTree (Hedges et al. 2006).

As a second biological example, we examined the utility of TraitRELAX in detecting changes in the selective type upon transitioning in the lifestyle of bacteria. To this end, 68 genes from 50 species of γ-proteobacteria, of which 36 are free living and 14 are endosymbionts were examined. The coding sequences, the phylogeny, and character state assignments of extant taxa (free living = 0, endosymbiont = 1) were retrieved from Husník et al. (2011). The sequences were aligned with RevTrans version 2.0b (Wernersson and Pedersen 2003) with default parameters. Positions with more than 90% gaps across sequences were filtered out.

In all analyses, branch lengths of the phylogeny were optimized using the M3 codon model (Yang and Nielsen 2002) as implemented in Bio++ (Guéguen et al. 2013). Comparisons between the null and alternative TraitRELAX models were conducted using the parametric bootstrap approach as detailed above.

### Availability

TraitRELAX was implemented as an open-source program in Bio++ (Guéguen et al. 2013) and its code is available at https://github.com/halabikeren/TraitRELAX. The input to the program is a tree, as well as sequence and character data files. The program outputs the maximum-likelihood score for the null and alternative models as well as their inferred model parameters.

## Results

### Using TraitRELAX to infer associations between selection intensity and phenotypes

In this work we developed the TraitRELAX model, which enables the detection of associations between the evolution of phenotypic traits and the intensity of selective forces operating on coding sequences. The model emulates the joint evolution of a binary phenotypic trait and the coding sequences using two Markov processes. Character evolution is described by a two-state Markov process, while coding sequence evolution is modeled using a variant of the branch-site model RELAX (Wertheim et al. 2015). In this joint model, each position in a coding sequence is subject to a selective regime (negative, neutral, or positive) whose intensity could increase (or decrease) upon a change in the phenotype. Namely, sequence evolution in segments of the phylogeny evolving under state ‘0’ (background, *BG)* is described by a set of selection parameters: *ω*_0_, *ω*_1_ and *ω*_2_, while segments evolving under state ‘1’ are mapped to the foreground *(FG)* category. Shift in the selection intensity upon character transition is modeled using a selection intensity parameter *k* that modifies the selection parameters on the *FG* segments (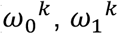 and 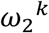), such that *k* < 1 indicates relaxation of selection and *k* > 1 indicates intensification. A likelihood ratio test examines whether the coding sequences are subject to a trait-dependent shift in the selection intensity (i.e., whether *k* is significantly different from 1). The full details of this model and likelihood computations are given in “Materials and Methods”. In the following, we first evaluate the performance of the method on simulated data sets. We next apply it on real data to detect changes in the selection intensity in the primate SEMG2 gene upon transition to a polygynandrous mating system and in 68 γ-proteobacteria genes upon transition to endosymbiotic lifestyle.

### Performance in simulations

Simulations were used to investigate the performance of TraitRELAX with regard to the false positive rate (FPR; i.e., detecting association between trait evolution and sequence evolution when no such association exists), sensitivity, and accuracy of parameter estimation.

### False positive rate and sensitivity

We analyzed the FPR by setting the selection intensity parameter *k* to 1, thereby simulating sequence data with no change in selection intensity along the phylogeny, and regarded as false positive any inference in which the null hypothesis (no association) is rejected according to the LR statistic. First, we examined the adequacy of p-values computed using the 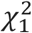 approximation to the LR distribution under null conditions. Similar to the analysis of Levy Karin et al. (2017), we found deviations of the FPR from the expected 5%. For example, for alignments with 300 codons and a significant threshold of *α* = 0.05 (5% expected FPR), the FPR was 16.5%, 21% and 15.5%, for simulations with 16, 32, and 64 taxa, respectively (Simulation Set 4). For this reason, we empirically determined a threshold for the LR statistic using parametric bootstrapping, such that the FPR is fixed at 5% (see Materials and Methods). The thresholds determined this way for simulations with 300 codon positions were 6.57, 7.62, and 6.79, for 16, 32, and 64 taxa, respectively (compared with 3.84 using the 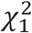 approximation). This procedure was performed for each combination of simulated parameters in the sensitivity analysis detailed below.

We analyzed the sensitivity of TraitRELAX based on simulations with various number of species, sequence lengths, and different values of the selection intensity parameter *k.* Our results indicated that the sensitivity of the method increases with data availability. This is true both for the number of sequence positions (**Error! Reference source not found.**2a) and the number of analyzed taxa (**Error! Reference source not found.**2b). For example, for simulations with 32 taxa and *k* = 0.5, TraitRELAX correctly rejected the null hypothesis in 46% of the simulations with 300 sequence positions and in 72% of the simulations with 600 positions. These numbers increased to 74% and 88% when using trees with 64 taxa. As may be expected, increasing the magnitude of the selection intensity parameter also resulted in increased sensitivity (e.g., 74% correct rejection in simulations under *k* = 0.5 and 84% in simulations under *k* = 0.2; Fig. 2b). Usage of the empirical threshold instead of the theoretical one (derived from the 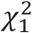 distribution) had little effect on the sensitivity of the method (Supp. Fig. 1). We next examined the effect of the number of character transitions by computing the sensitivity in simulations with increasing values of the character transition rate, *μ.* In these simulations, a bell-shape pattern was observed, where maximum sensitivity was obtained for intermediate values of *μ* and dropped when very small or very large number of character transitions were simulated (**Error! Reference source not found.**2c). In comparison, we also optimized the sequence model with branch partitions that were derived from the true (simulated) character histories, thus serving as a reference for the optimal performance that could be expected when eliminating any error in the reconstructed character history. As expected, the sensitivity in this case was consistently higher. The difference between using the true history rather than inferring it, was particularly apparent for high values of *μ,* representing simulations with a large number of simulated character transitions (Fig. 2c). Evidently, the complex character histories in such cases are harder to infer and reduce the ability of the method to detect genuine trait-related shifts in the selection intensity.

**Figure 2.**
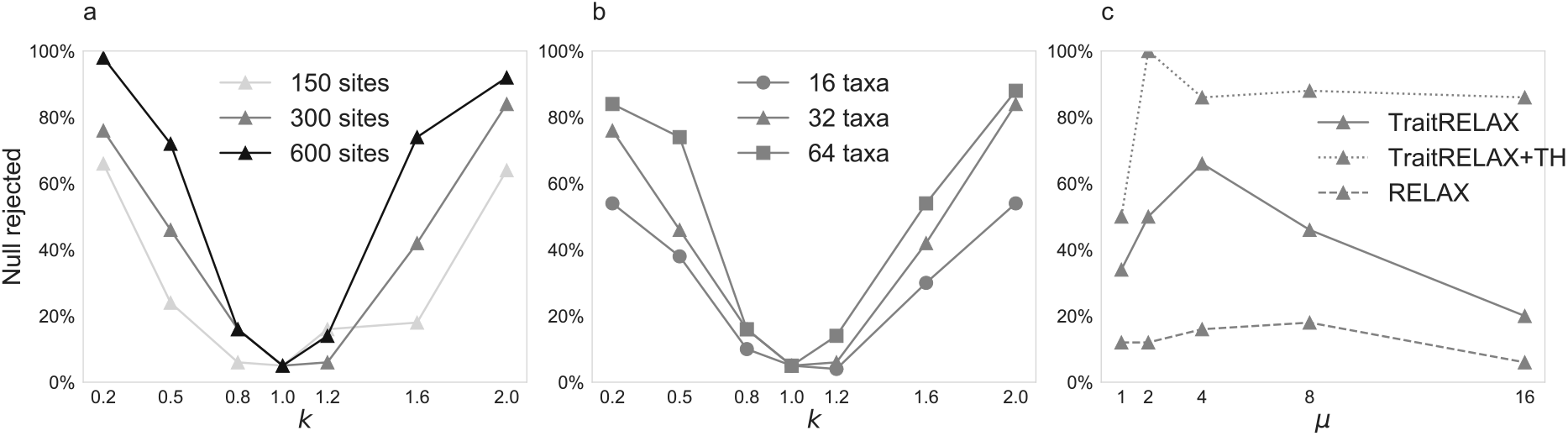
Assessment of FPR and sensitivity based on simulations with different data sizes. The percent of replicates for which the null model was rejected is shown for simulations with (a) 32 taxa and different number of codon positions, and (b) 300 positions and different number of taxa. (c) Sensitivity is shown for different values of the simulated *μ* parameter. Results are shown for standard TraitRELAX (solid line), TraitRELAX given the true character history (TH) and sequence optimization only (dotted line) and RELAX given a maximum parsimony partition (dashed line). In all TraitRELAX cases, parametric bootstrap was used to determine an empirical LR cutoff for rejecting the null hypothesis. When using RELAX, the LR threshold was based on the 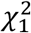. For both methods the cutoff for rejection was set at *α* = 0.05.

### Accuracy of parameter estimation

We examined the accuracy of the method in estimating the selection intensity parameter *k,* as a function of the number of taxa and codon positions. In the null case of *k* = 1, the average inferred *k* was close to the simulated value, regardless of the number of taxa (Fig. 3a) or codon positions (Supp. Fig. 2d-f). In other cases, the inferred value of *k* was conservative with respect to the magnitude of the change in the intensity of selection, such that it was generally higher than the true value when relaxation of selection was simulated (*k* = 0.2) and lower than the true value when intensification was simulated (*k* = 2). This bias was most pronounced when only few taxa were available and nearly disappeared for the case of 64 taxa and 600 positions (Fig. 3a, Supp. Fig. 2a,c,e). Similar to the sensitivity analysis, we found that the averaged error in the inferred value of *k* is most pronounced when the number of character transitions is very high or very low, and lowest when intermediate values of *μ* were simulated (Fig. 3b).

**Figure 3.**
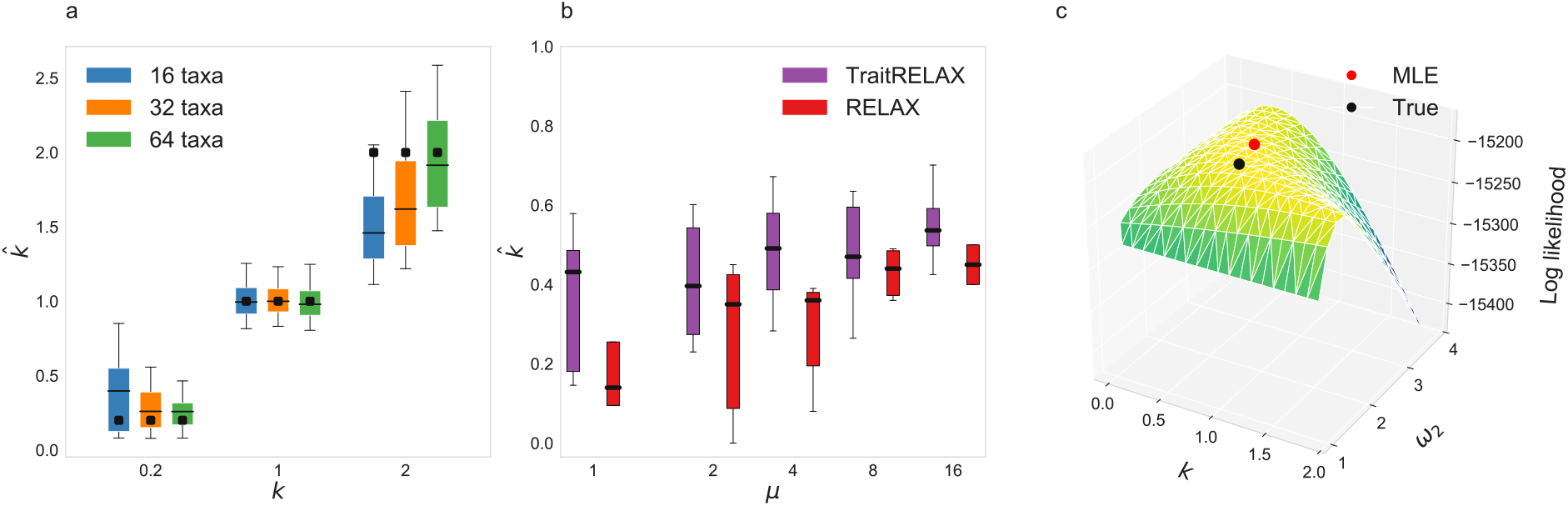
Accuracy of inferring the selection intensity parameter. (a) The distribution of the inferred values of *k* for simulations with 600 positions. For each simulated *k* value, the numbers of taxa are ordered from left to right: 16, 32, and 64 taxa. The horizontal lines within the box plots indicate the mean inferred values and the simulated values are shown as black dots. (b) The inferred values of *k* are shown for simulations with 32 taxa, 300 positions, and *k* = 0.5 for different values of the simulated *μ* parameter. (c) Log likelihood surface over a grid of 300 combinations of *ω*_2_ and *k.* The panel depicts a single simulation instance in which *k* = 0.8, *ω*_2_ = 2 (with *p*_2_ = 0.99) with 32 species and 600 codon positions. The simulated values and maximum likelihood estimators are shown in black and red dots.

Finally, we studied the interplay between the inference of *k* and that of the **ω** parameters. The joint examination of the inferred values of *k* and the **ω** parameters suggests that underestimation of *k* (in the case of intensification of selection) is compensated by overestimation of *ω_2_* values, while the opposite is observed in the case of relaxation (i.e., overestimation of *k* and underestimation of *ω*_2_) (Supp. Fig. 3). Indeed, examining the joint likelihood surface of *k* and *ω*_2_ revealed that both the true (i.e., simulated) values and their maximum likelihood estimates are located along a ridge of high likelihood values, such that several (*k, ω*_2_) combinations receive similar log-likelihood scores (Fig. 3c), leading to difficulties in inferring the exact value of each parameter separately.

### Comparison to RELAX

The simulated data were used to compare the performance of TraitRELAX to that of the branch-site model RELAX (Wertheim et al. 2015), as implemented in HyPhy (Kosakovsky Pond et al. 2005). To execute RELAX, a partition of the input phylogeny to background and foreground branches should be provided, corresponding to putative branches evolving under character state ‘0’ and ‘1’, respectively. To this end, we used the DELTRAN approach (Farris 1970; Swofford and Maddison 1987), implemented in Bio++, to infer the maximum parsimony solution given the character states of the extant taxa in each simulated data set.

Similar to the results obtained by TraitRELAX (Fig. 2), the sensitivity of RELAX increased with the intensity of the simulated value of *k* and with the size of the data (Supp. Fig. 4). Here too, RELAX had higher sensitivity in simulations with intermediate values of the character transition rate parameter *μ,* although the trend was shallower compared to that observed for TraitRELAX (Fig. 2c). Noticeably, the sensitivity of RELAX was consistently lower than that of TraitRELAX across all simulation scenarios (Fig. 2c, Supp. Fig. 4). For example, in simulations with 64 taxa, 600 positions, and *k* = 0.2, TraitRELAX correctly rejected the null in 98% of the cases, while RELAX in 44%.

Next, we evaluated the accuracy of both methods in inferring *k* in simulations with varying data size and with different number of character transitions. In all simulated conditions, the error and bias of RELAX was consistently higher than that of TraitRELAX (Fig. 3a-b, Supp. Fig. 5). Taken together, these analyses suggest that by integrating character trait data into a joint statistical framework, TraitRELAX is more robust than RELAX to uncertainty in trait evolution.

### Association between selection type and intensity upon transition to polygynandry in primates and to endosymbiosis in γ-proteobacteria

We demonstrate the use of TraitRELAX to detect trait-related changes in selection intensity using two biological data sets: one concerning transitions in mating systems among primates, and the other between free living and endosymbiont bacteria. Transition among mating systems in primates has been previously associated with shifts in the selection patterns operating on several reproductive genes, suggesting that sperm competition among males in polygynandrous societies, in which multiple males complete for the same female, is an underlying force of sexual selection (Ramm et al. 2008). Elevated nucleotide substitution rate and *dN/dS* ratio in polygynandrous lineages has been detected in the Semenogelin II (SEMG2) gene, whose encoded protein takes part in the formation of a gel matrix that encases ejaculated spermatozoa (Dorus et al. 2004; Hurle et al. 2007; O’Connor and Mundy 2009; Levy Karin et al. 2017). Here, we used TraitRELAX to investigate associations between transitions in primates mating systems and shifts intensity of selection in SEMG2. The SEMG2 gene coding sequences of 24 primate species were collected (see Material and Methods), nine of which are polygynandrous and 15 that participate in other mating systems (including monogamous, polygamous, and polygamous systems). The distribution of mating systems along the phylogeny indicated that there were at least six transitions among this group of analyzed species (based on the maximum parsimony reconstruction, which can be considered as lower bound). A comparison between the null and alternative models of TraitRELAX revealed significant intensification of selection under the polygynandrous state (*k* = 2.13, p-value = 0.03). We found that most of the SEMG2 sites are dominated by neutral selection and are not affected by transitions in the mating system 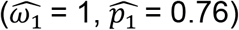, while approximately 17% of the sites are subject to positive selection, which is intensified along the polygynandrous lineages (with *ω*_2_ = 2.33 and 6.08 in the ‘other’ and polygynandrous states, respectively). This proportion is in agreement with a previous study that detected positive selection in the SEMG2 gene with a proportion 0.15 of sites and *ω*_2_ = 3.27 (Ramm et al. 2008).

Next, we investigated whether transitions in the lifestyle of γ-proteobacteria are associated with shifts in the selection intensity operating on 68 housekeeping genes. Unlike free-living bacteria, bacterial endosymbionts rely on their host for mandatory functions, such as translation and metabolite decomposition (Husník et al. 2011). The smaller population size of the endosymbionts, caused by their inability to replicate independently from the host, reduces the efficiency of natural selection and can result in accumulation of deleterious mutations and overall elevated rates of nonsynonymous substitutions and radical amino acid replacements (Muller 1964; Moran 1996; Funk et al. 2001; Herbeck et al. 2003; Wernegreen 2011). Indeed, a relaxation of selection in endosymbiotic bacteria, relative to their free-living progenitors, was detected using the RELAX model (Wertheim et al. 2015). However, that analysis was performed on an alignment of 69 single-copy genes that were treated as a single concatenated gene region and thus did not provide information concerning the identity of specific genes that are subject to differential selection in endosymbionts and the magnitude by which they are affected. Here, we used TraitRELAX to analyze 68 of these genes separately (one gene was excluded from analysis due to the sparse sequence data). The analysis was performed on 50 species of γ-proteobacteria (16 endosymbionts – state 1, and 34 free living – state 0), with at least seven transitions between the two states according to maximum parsimony reconstruction. Comparing the null and alternative TraitRELAX models revealed significant change in selection intensity under endosymbiotic lifestyle in sixteen out of the 68 examined genes (*p-value* < 0.05 following the Benjamini-Hochberg false discovery rate correction for multiple testing (Benjamini and Hochberg 1995), with relaxation of selection in ten genes (with 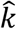 ranging from 0.78 to 0.94) and intensification in six genes (with 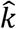 ranging from 1.06 to 1.36). The relaxed genes were either involved in metabolite decomposition or in tRNA selection processes, while the intensified genes were involved in transcription and translation (Supp. Table 1). In all cases, the inferred value of *ω*_2_ was 1, suggesting that in either state positive selection has not operated on these genes and the change in selection intensity is with respect to purifying selection only. For the ten relaxed genes, the mean estimated values of *ω*_0_ and *ω*_1_ were 0.016 and 0.140 in the free-living state and 0.026 and 0.178 in the endosymbiotic state, respectively and for the six intensified genes the mean values of *ω*_0_ and *ω*_1_ were 0.013 and 0.137 in the free-living state and 0.006 and 0.097 in the endosymbiotic state.

## Discussion

In this study we developed TraitRELAX, a new method for detecting associations between organismal traits and selection intensity at the codon level. Within the framework of the method, an association is expressed as a dependency between phenotypic trait evolution and changes in patterns of selection operating on coding sequences. An important consideration of the method is that transitions in the phenotypic states should affect all sequence positions simultaneously, and thus the evolution of each codon site should not be treated independently. Since the exact timings of phenotypic transitions are unknown, we consider possible scenarios that led to the observed phenotypic states at the extant taxa by using stochastic mappings, such that each mapping induces a possible partition as input for the sequence model and all mappings are considered simultaneously. This, however, entails heavy computations since likelihood computation of the sequence data is performed multiple times for each parameter combination examined.

To speed up the likelihood computation, we adopted a similar approach to that presented in Levy Karin et al. (2017), in which multiple stochastic mappings are represented by a single expected history (Equation 10). Here, unlike in their paper, all possible mappings were summarized analytically using the rewards method (Minin and Suchard 2008). This approximation limits the number of transitions along each branch to at most two (see Supp. Text 1). When many transitions have occurred along a branch, this approach is expected to yield inferior approximation to the exhaustive approach that uses multiple stochastic mappings (Equation 9). However, most organismal traits of interest are key features of a lineage and are expected to evolve rather slowly, such that multiple or back transitions on the same branch are expected to be rare. In such cases, the expected history approximation should perform well. In addition to reducing running times, the approximation approach increases the stability of the computation as it is robust to the stochasticity of the sampled mappings. Another alternative approximation can be obtained by clustering the set of stochastic mappings into several groups and perform the computation on a representative of each, weighted by its relative frequency. Such an approximation was adopted by Jones et al. (in press). The method groups stochastic mappings in case phenotype transitions have occurred on the same branches and ancestral state assignments are identical. While allowing for multiple unique phenotype histories, this approach practically restricts phenotype transitions to occur at internal nodes and limits the number of transitions to at most one per branch.

TraitRELAX can receive as input an ultrametric tree, in which branch lengths are proportional to time. Such trees well describe traits whose rate of change is time-dependent, such as organismal habitat or lifestyle. Alternatively, a non-ultrametric tree can be provided, better reflecting traits whose rate of change is proportional to the amount of genetic change or those whose transition rate is likely associated with the generation time. Because the reconstruction of time-calibrated phylogenies suffers from high degrees of uncertainty (Dornburg et al. 2012; Wertheim et al. 2012; Dos Reis et al. 2015; Bromham et al. 2018), one may prefer the use of uncalibrated trees, unless there is a strong evidence that the evolution of the trait in question is time dependent.

Our simulation study demonstrated that the sensitivity and accuracy of the method depend on the size of the data, the magnitude of selection-intensity shift upon phenotype transition, and the number of character trait transitions. When there are only few character state transitions across the phylogeny, there is little information to accurately infer changes in selection intensity. When the character history is too complex (high *μ* values), there is not enough information in the character data of extant species to accurately reconstruct the character history, and thus, reducing the ability to detect phenotype effect on patterns of selection at the sequence level. Notably, in small data sets, failure to reject the null hypothesis could indicate no association between the examined trait and coding sequence evolution, but could also indicate low statistical power to detect such an association, especially if the inferred value of the selection intensity parameter *k* is far from 1. To test whether the data are sufficient for reliable inference, sensitivity analysis should be conducted (Boettiger et al. 2012). To this end, simulations should be used to generate artificial data sets with similar data characteristics as the empirical data (in terms of number of taxa, sequence sites, and number of character transitions), and with various values of the *k* parameter that are increasingly distant from 1. Such artificial data sets, as generated in this study, can guide us on the possible values of *k* for which the null hypothesis could be reliably rejected.

When compared to the existing branch-site model RELAX (Wertheim et al. 2015), TraitRELAX demonstrated higher sensitivity to detect trait-related changes in selection intensity and higher accuracy in estimating the *k* parameter governing the direction and magnitude of the selection shift. Multiple differences between the methods can explain the observed results. First, unlike RELAX, TraitRELAX considers the uncertainty in the trait history and additionally allows trait transitions to occur anywhere along a branch rather than only at hypothesized speciation events (i.e., internal nodes of the phylogeny). Second, in TraitRELAX we constrain the selective regimes assigned to each position to remain the same across the phylogeny, while the implementation of RELAX in HyPhy allows changes in the selective regime of a site (e.g., from purifying to positive selection) when transitioning across branches (for more details on the difference between these implementations see Supp. Text 3). This latter approach could result in confounding effects between the selection intensity parameter *k* and the *ω* parameters and could lead to erroneous estimates of *k* (Supp. Fig. 5a). For example, when the inferred value of *k* is very small, 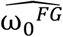 may resemble 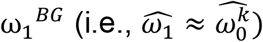. In such a case, the null scenario in which a sequence position evolves under neutral selection throughout the entire phylogeny yields similar likelihood to an alternative scenario in which the same position shifts from neutral selection in the *BG* branches to purifying selection in the *FG* branches. Finally, we note that the sensitivity and accuracy of RELAX observed here were lower than those reported by Wertheim et al. (2015), most probably due to the smaller data size examined in our simulations (reaching up to 600 codon positions, compared to 21,154 positions in theirs).

A main benefit of the statistical framework presented here is that it simultaneously emulates the evolution of both the examined phenotype and that of the coding sequences. As such, it deviates from many widely used models that rely on a two-step inference scheme, in which the evolution of the phenotype is first reconstructed, and this inference is then incorporated into models of sequence evolution while ignoring any uncertainty in the preceding inference step. The TraitRELAX joint phenotype-sequence approach can be extended in various ways, most notably, by selecting a different branch-site codon model as the underlying sequence model. For example, a generalization of our approach can be applied by mapping independent codon models (Yang et al. 2000) to each phenotypic state, while having some of the parameters shared between the models to reduce the overall number of parameters. This will enable the detection of more subtle associations between the evolution of the phenotype and that of the examined sequences. One such example was recently presented by Jones et al. (in press) who developed the PG-BSM method that, in common with the approach presented here, integrates trait and coding sequence data in a joint statistical framework. The PG-BSM method is designed to detect phenotype-dependent shifts in adaptation, that is, persistent or transient increase of *ω* at a site dictated by the trait under study. The PG-BSM method further allows for a proportion of the sequence positions to evolve in a trait-independent manner. Incorporating such a parameter has been proved useful in other models (Levy Karin et al. 2017), and thus may be a beneficial extension to TraitRELAX. Lastly, a useful extension of TraitRELAX would allow consideration of categorical phenotypes that consist of more than two states, which will also allow the examination of multiple, perhaps correlated, traits. The combination of multiple traits can be perceived as a single, categorical, trait, whose states correspond to the various combinations of character states of the traits of interest. Such modelling would provide more realistic scenarios regarding the many pathways by which a combination of organismal phenotypes governs evolutionary processes at the genomic level.

## Acknowledgments

The authors thank Prof. Sergei L. Kosakovsky Pond for his useful advice during the development process of the method and Julien Y. Dutheil for his assistance in the implementation of the method.

