## Supplementary text for "A codon model for associating phenotypic traits with altered selective patterns of sequence evolution"

[4] Swedish Collegium for Advanced Study. Thunbergsvägen 2
752 38 Uppsala, Sweden.

* To whom correspondence should be addressed:

Running title: Coding sequence - phenotype integrated model

Keywords: Evolutionary selection; intensification, relaxation, genotype-phenotype; γ-proteobacteria; SEMG2;

### Supplementary Text

### Supplementary Text 1

##### The expected history approximation

The likelihood of the sequence data is computed while conditioning on the history of trait transitions. Here we provide more details on how this computation is performed. Let $H$ denote a random variable representing the history of character state transitions along the phylogeny $T$, and let $h$ be a single possible realization of $H$. We define a new tree, $T_h$, where each point of a trait transition defines an internal node with a single descendent lineage.

The expected history, $E(h)$, is a single history that summarizes a set of possible trait histories. The expected history is constructed by computing the expected time spent in each state in each branch over the entire set of histories. These can be computed in two alternative approaches: a sampling-based approach, in which these values are obtained as the average over a finite sample of $N$ stochastic mappings (as done in Mayrose and Otto 2011; Levy Karin et al. 2017), or in an analytic approach, which considers all possible trait histories and uses the rewards method of Minin and Suchard (2008). As described below, in our implementation we followed the more accurate analytic computation.

First, the ancestral states are determined based on the posterior probabilities of character states at internal nodes, as obtained from the character model and data (Nielsen 2002). Let $f_{n,i}$ be the posterior probability of observing character state $i$ in ancestral node $n$. Then, the state assignment to node $n$ in $E(h)$ is $argmax_i(f_{n,i})$.

Next, the expected times spent in character states ‘0’ and ‘1’ and in each branch $b$ along $T$, termed here $E\left(t_0^b\right)$ and $E\left(t_1^b\right)$, are computed. Specifically, we compute for each branch in the phylogeny the total evolutionary reward for the expected times spent in the two character states. This value is computed based on a fixed set of rewards $\{\omega_0, \omega_1\}$ for character states ‘0’ and ‘1’, respectively (equations 1.4-2.12 in Minin and Suchard 2008). As such, setting $\omega_0$ to 1 and $\omega_1$ to 0 yields a total evolutionary reward value that is equal to $E\left(t_0^b\right)$. Similarly, $E\left(t_1^b\right)$ can be obtained by setting $\omega_0$ to 0 and $\omega_1$ to 1.

Once the above expectations are computed, the location of state transition along each branch (if there was one) has to be resolved. Consider a branch, $b$, whose parent node is assigned with state '1' and child node with state '0' (or vice versa) and the expected time spent in state '1' is $E\left(t_1^b\right)$. In this case, the expected history is defined to contain a single transition, whose location is $E\left(t_1^b\right)$ time units away from the parent node. Alternatively, in case both parent and child nodes are assigned with the same state there must be at least two transitions along the respective branch (if neither $E\left(t_0^b\right)$ nor $E\left(t_1^b\right)$ is equal 0). In this case, the time spent in the shared state is divided such that there are two transitions along the corresponding branch (from the shared state, to the alternative state, and then back to the shared state). The locations of state transitions are determined based on the posterior probabilities of the shared state at the two terminal nodes. Specifically, let $b$ be a branch of length $t$ in which the inferred state at both parent and child nodes is ‘0’, the expected time spent in state ‘0’ is $E\left(t_0^b\right)$, and the expected time spent in state ‘1’ is $t - E\left(t_0^b\right)$. Then, *b* is divided to three segments, such that the middle segment (in red below) is assigned to state ‘1’ and the two terminal segments (in black) to state ‘0’. The lengths of the segments are determined as follows:

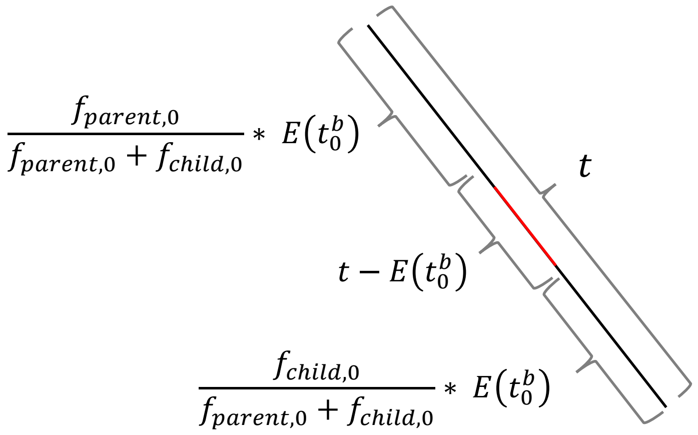


where $f_{parent,0}$ and $f_{child,0}$ are the posterior probabilities of being in state ‘0’ at the father and child nodes, respectively.

### Supplementary Text 2

##### Adequacy of the approximated likelihood function

The full likelihood function of TraitRELAX (Equation 6 in the main text) integrates over all possible character histories. Since this integration is not feasible, the computation is approximated by a set of $N$ sampled stochastic mappings (Equation 9 in the main text denoted here the exhaustive approximation). The accuracy of this approximation increases with the number of sampled histories, $N$, but becomes prohibitively long even for a modest value of $N$, given the large size of the codon rate matrix. When examining its performance, we chose $N$ = 1,000 as a balanced tradeoff of accuracy/running time. Alternatively, this computation is approximated by a single expected history (see Supplementary Text 1 and Equation 10 in the main text) which is computed analytically (denoted here as the expected history approximation). While this heuristic computation yields markedly lower running times, it could result in inaccurate likelihood computations. Thus, we examined the adequacy of these approximations by measuring the difference in the computed log-likelihood values compared to that obtained using the true, simulated, history.

As shown in Supplementary text 2, Fig. 1, the log-likelihoods obtained using the expected history approximation were consistently more similar to those of the true history as compared to those obtained using the exhaustive approximation. Moreover, the variance of the log-likelihood was greater when using the exhaustive approximation than when using the expected history approximation. Additionally, the expected history approximation is deterministic and thus yields a stable likelihood computation, unlike the exhaustive approximation which is susceptible to the stochasticity of the sampled mappings. Together, this suggests that the expected history approximation is both closer to the true history as well as more robust and we therefore opted to use this approximation in all analyses conducted in this study.

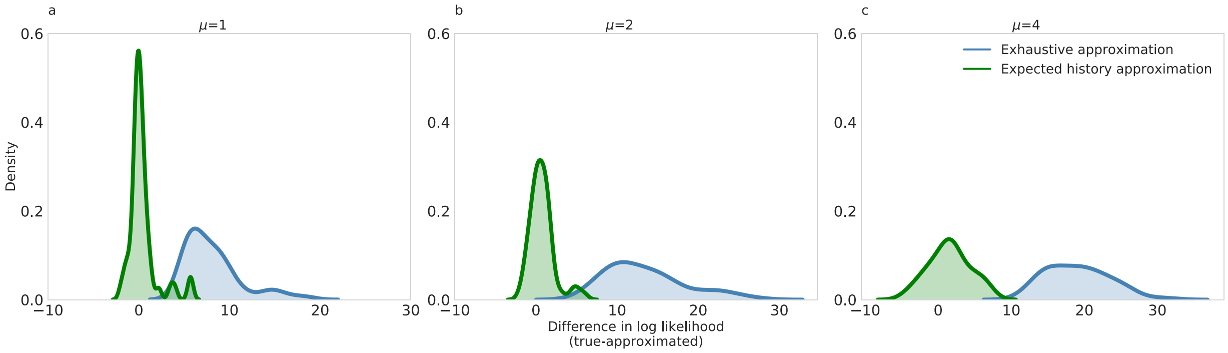


***Supplementary Text 2, Figure 1. Comparison of the log-likelihoods obtained using the exhaustive and expected history approximations.*** *The distribution of the differences between the log-likelihood values obtain using the true (simulated) history and those obtained using either the expected history approximation (green) and the exhaustive approximation (blue). The distributions are computed for 50 simulations of trees with 32 taxa, 300 codon positions and* $k$ *= 0.5 with character transition rate of (a)* $\mu$ *= 1, (b)* $\mu$ *= 2, and (c)* $\mu$ *= 4.*

### Supplementary Text 3

#### Sequence likelihood computation approach

The sequence model of TraitRELAX consists of three site categories that model the different selective regimes that operate on the codon positions using three $\omega$ parameters ($\omega_0$, $\omega_1, \omega_2$ for purifying, neutral and positive selection) and two branch categories ($BG$ and $FG$ corresponding to segments evolving under state '0' and '1', respectively) that model changes in the intensity of selection in the $FG$ branches relative with the $BG$ branches using a selection intensity parameter $k$. As such, each selective regime $c$ is modeled in the $BG$ branches by the value of the respective $\omega_c$ parameter and by the value $\omega_c^k$ in the $FG$ branches.

An important, yet overlooked, consideration when computing the likelihood of a branch-site codon model concerns with the nature of shifts between site categories (in this case, selective regimes) across branches. Consider three adjacent branches, such that $b_1$ and $b_2$ are the descendant lineages of $b_3$. There are three ways to compute the sequence likelihood when moving across branches. The first option, which we followed in the implementation of TraitRELAX, restricts the site category in all branches to be the same, whether they belong to the same branch category or not. Thus, if a position evolved under $\omega_0$ in $b_3$, it would evolve under $\omega_0$ also in $b_1$ and $b_2$. Notably, because each branch category has its own value of $\omega$ for each site category (e.g., $\omega_0$ of the $FG$ category is set to $\omega_0$ of the $BG$ category raised to the power of $k$) the site can still experience a change in the magnitude of selective pressure upon transitioning from $b_3$ to $b_2$, but not when transitioning between two adjacent branches that belong to the same branch category.

In a second alternative, the site category assigned to each position may vary between adjacent branches only if the ancestral one is assigned to the $FG$ category and its descendent is assigned to the $BG$ category. In this case, a site evolving under $\omega_0$ in $b_1$, may evolve under any $\omega$ category in $b_2$. A possible shortcoming of this approach is that it allows for more combinations of site-categories assignments in certain branch partitions. Specifically, if a transition between branch categories occurs in an internal branch (e.g., $b_3$ is assigned as $FG$ while $b_1$ and $b_2$ as $BG$), the constraint on the $\omega$ assignment for the two descendant branches is relaxed, such that a site may evolve under $\omega_0$ in $b_3$, $\omega_1$ in $b_2$, and $\omega_2$ in $b_1$. On the other hand, if the transition between branch categories occurs in an external branch (e.g., $b_1$ is assigned as $FG$ while $b_3$ and $b_2$ as $BG$), the constraint on the $\omega$ assignments is relaxed only for the single external branch ($b_1$). This issue could bias multiple testing procedures that iteratively assign a single branch to the $FG$ category and test for the presence of positive selection on that branch (e.g., Anisimova and Yang 2007) as it might favor certain partitions of branches that allow for more combinations of assignments to site-categories. Because this approach is the one most commonly used by the community (for example, in the popular software package PAML; Yang 1997), further research is needed to explore its behavior.

In a third and most permissive approach, which is used for example by Wertheim et al. (2015) and implemented in the HyPhy package, the site category assigned to each position may vary across all branches, regardless of their assigned branch category. This approach allows for rapid shifts in site categories that are not necessarily realistic, such as shifts from purifying to positive selection along very short branches, and can thus lead to a high rate of false positives (Kosakovsky Pond and Frost 2005). In addition, this approach could lead to confounding effects between parameters in the model (e.g., between the selection intensity parameter $k$ and the $\omega$ parameters; see Discussion in the main text).

### Supplementary Figures

#### Supplementary Figure 1

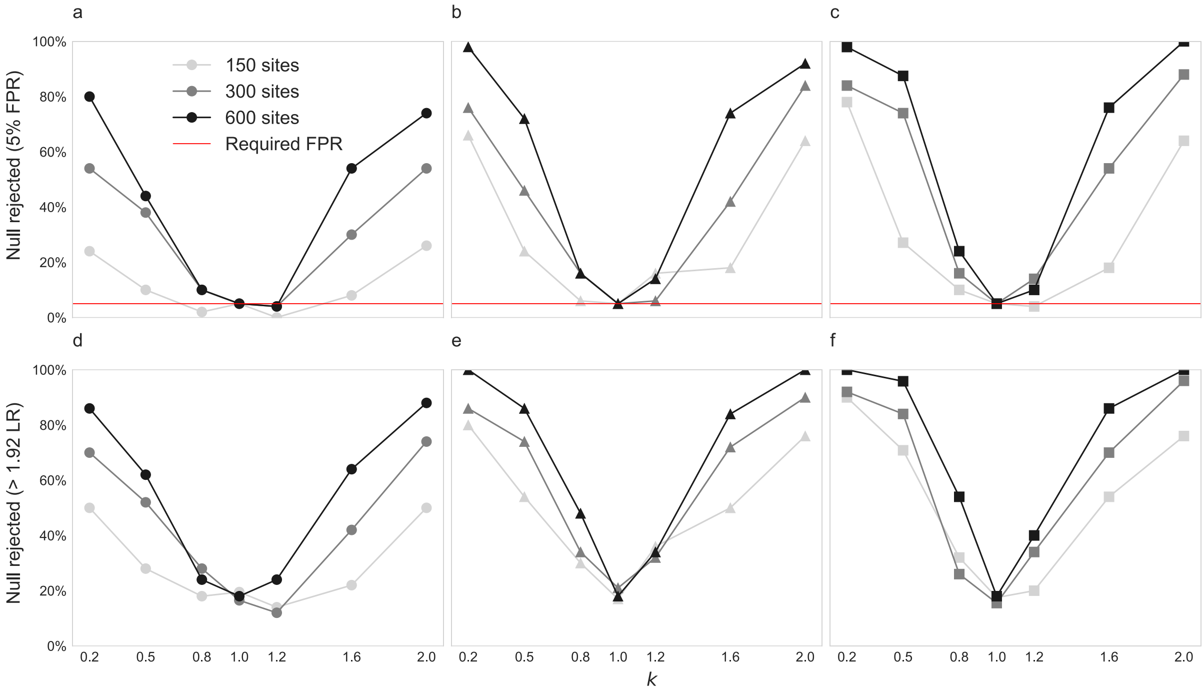


**Sensitivity and FPR assessment for various data sizes.**

The percent of simulated datasets in which the null hypothesis was rejected by TraitRELAX using the LRT when the LR threshold was determined according to parametric bootstrapping (top row; a-c) or approximated using the $\chi^2$ distribution (bottom row; d-f) with a significance threshold of $\alpha$ = 0.05. Results are shown for different number of taxa: 16 (panels a and d), 32 (b and e), and 64 (c and f).

#### Supplementary Figure 2

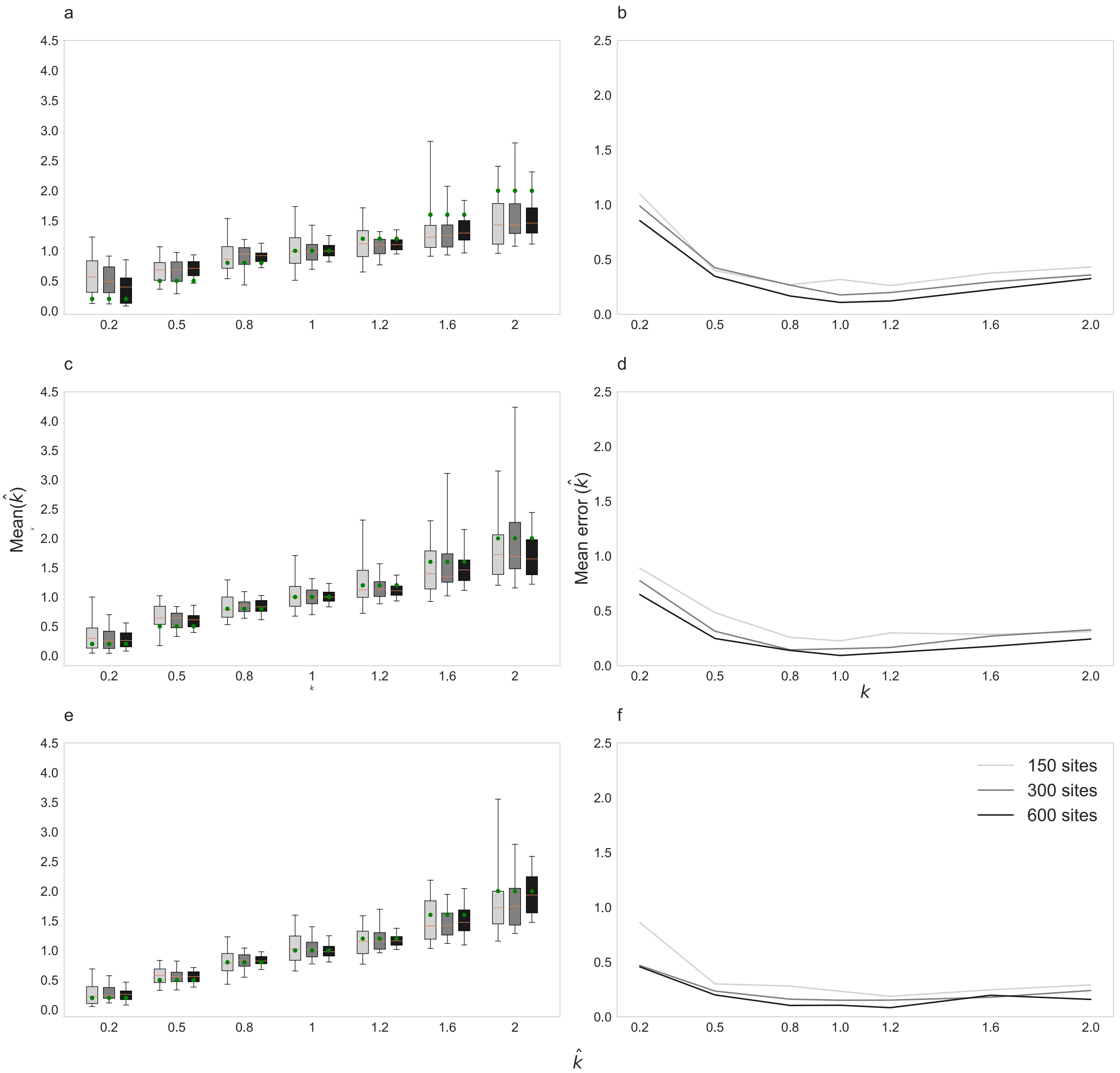


**Accuracy of inferring $k$ as a function of data sizes.** (a,c,e) The distribution of the inferred $k$ parameter by TraitRELAX and (b,d,f) its average inference error for simulations with different number of species: 16 (panels a-b), 32 (panels c-d) and 64 (panels e-f). The horizontal lines within the box plots indicate the median inferred values and the red dots indicate the simulated (true) values of $k$. The error is measured as $\left|log(\hat{k}) - log(k)\right|$ to account for the exponential effect of $k$ on the $\omega$ values of branch category 1.

**Supplementary Figure 3**

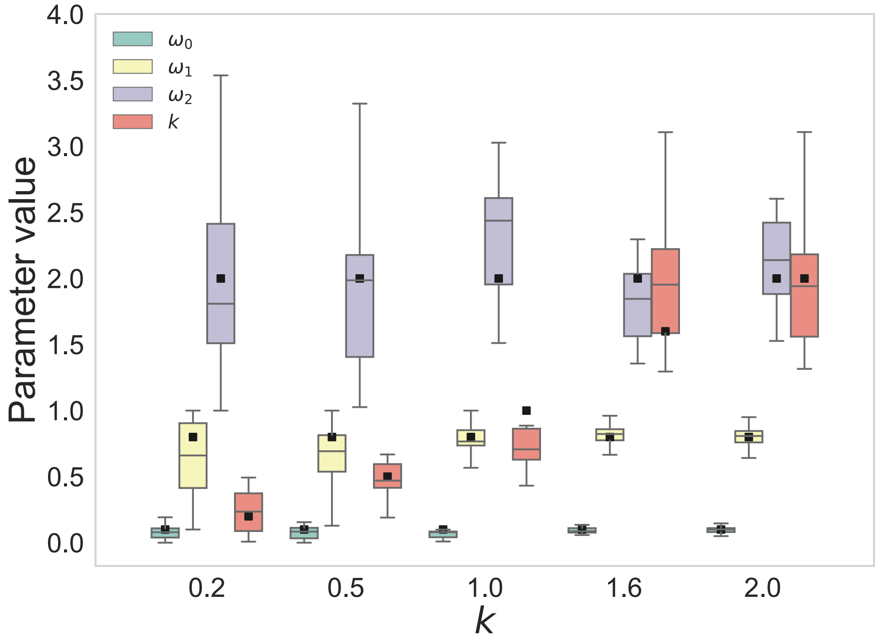


**The distribution of the inferred $k$ and $\omega$ parameters for simulations with various values of $k$.** For each simulated $k$ value, the box plots are ordered from left to right for the inference of: $\omega_0$, $\omega_1$ $\omega_2$, and $k$. The horizontal lines within the box plots indicate the median inferred values and the black dots indicate the simulated (true) values. All simulations were conducted with 32 taxa and 300 codon positions.

**Supplementary Figure 4**

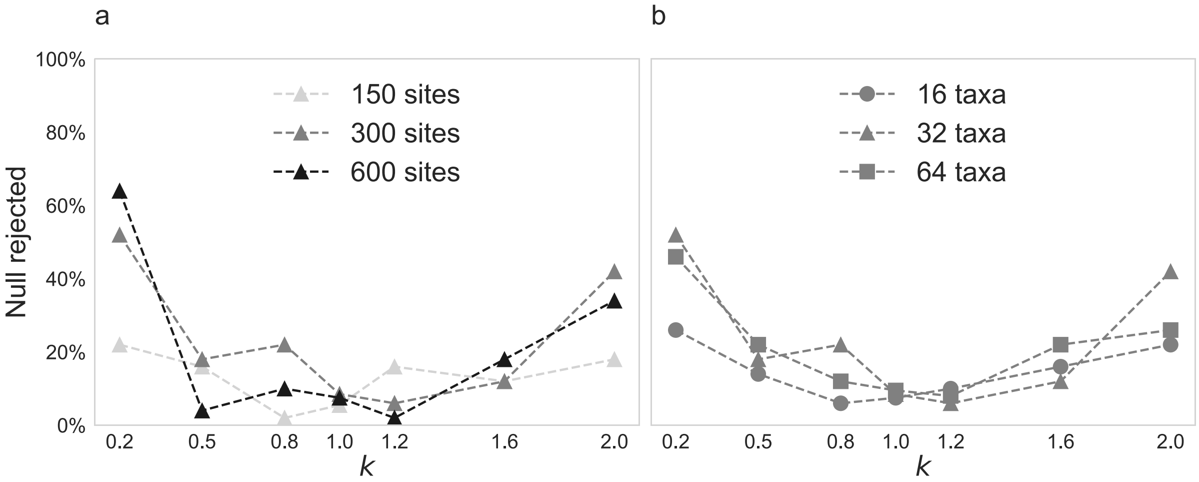


**Sensitivity assessment of the branch-site model RELAX**. The percent of replicates for which RELAX rejected the null model is shown for simulations with (a) 32 taxa and different numbers of codon positions, and (b) 300 codon positions and different numbers of taxa. Rejection of the null hypothesis was determined using the $\chi_1^2$ approximation to the LRT as recommended in Wertheim et al. (2015). In all cases, the cutoff for rejection was set at $\alpha$ = 0.05. In comparison to TraitRELAX, the sensitivity of RELAX was lower under all simulation conditions (see Fig. 2 in the main text).

#### Supplementary Figure 5

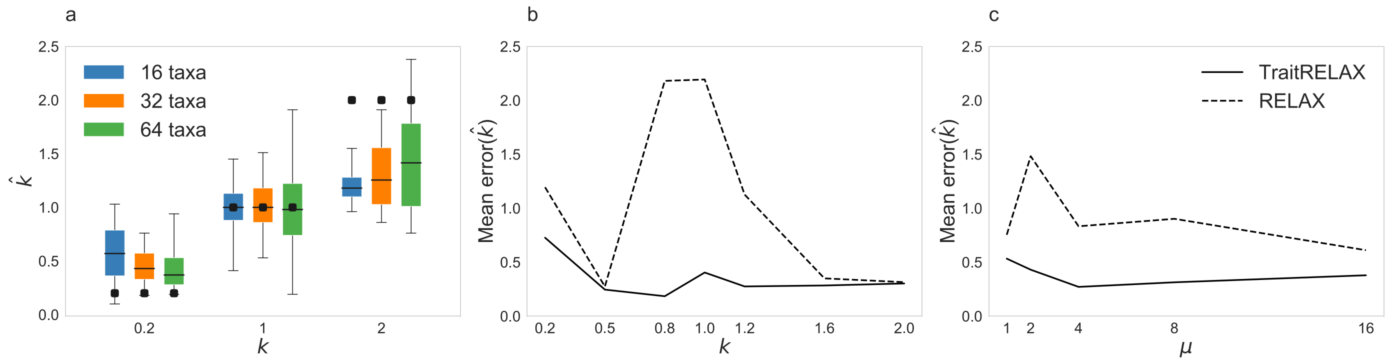


**Comparison of parameter estimation accuracy between TraitRELAX and RELAX**. (a) The distribution of the inferred values of $k$ by RELAX are shown for simulations with increasing number of taxa and 600 positions. The horizontal lines within the box plots indicate the median inferred values and the simulated values are shown as black dots. For comparison, the distribution of the inferred $k$ parameter by TraitRELAX is given in figure 3 of the main text. (b) The mean error of the inferred values of $k$ by TraitRELAX (solid lines) and RELAX (dashed lines) is shown for simulations with 32 taxa, 300 positions, and increasing values of $k$ (for a fixed value of $\mu$ = 8) and (c) with increasing values of $\mu$ (for a fixed value of $k$ = 0.5). The error is measured as $\left|log(\hat{k}) - log(k)\right|$ to account for the exponential effect of $k$ on the $\omega$ values of branch category 1.

**Supplementary Table 1. Inference of TraitRELAX on the 68 γ-proteobacteria house-keeping genes.** Genes for which a significant result was obtained following the false discovery rate (FDR) correction for multiple testing (Benjamini and Hochberg 1995) at $\alpha$ = 0.05 are marked with an asterisk. The $p$-values were computed based on the parametric bootstrapping procedure and then adjusted using the FDR procedure, yielding the $q$-values.

| Gene | Null log likelihood | Alternative log likelihood | Inferred model parameters | | | | | | |
|---|---|---|---|---|---|---|---|---|---|
| | | | $q$-value | $\omega_0$ | $\omega_1$ | $\omega_2$ | $p_0$ | $p_1$ | $k$ |
| | | | relaxed genes | | | | | | |
| rpsI | -7058.83 | -7020.19 | 0.02* | 0.00 | 0.10 | 1.00 | 0.50 | 0.44 | 0.78 |
| yeaZ | -23683.94 | -23668.87 | 0.02* | 0.03 | 0.21 | 1.00 | 0.45 | 0.44 | 0.84 |
| pgi | -41974.07 | -41951.28 | 0.02* | 0.01 | 0.14 | 1.00 | 0.62 | 0.32 | 0.84 |
| ybeY | -13392.69 | -13384.74 | 0.02* | 0.02 | 0.18 | 1.00 | 0.57 | 0.35 | 0.87 |
| asnS | -36378.68 | -36360.03 | 0.02* | 0.01 | 0.11 | 1.00 | 0.54 | 0.40 | 0.88 |
| mnmA | -29626.72 | -29614.07 | 0.02* | 0.01 | 0.11 | 1.00 | 0.53 | 0.43 | 0.91 |
| holB | -39217.16 | -39210.18 | 0.06 | 0.04 | 0.24 | 1.00 | 0.40 | 0.54 | 0.91 |
| argS | -48722.57 | -48704.89 | 0.02* | 0.02 | 0.15 | 1.00 | 0.59 | 0.36 | 0.91 |
| rpsQ | -4830.86 | -4829.68 | 0.77 | 0.01 | 0.13 | 1.00 | 0.44 | 0.53 | 0.91 |
| ftsY | -42810.06 | -42800.02 | 0.02* | 0.02 | 0.14 | 1.00 | 0.35 | 0.31 | 0.91 |
| fold | -23296.47 | -23290.92 | 0.06 | 0.01 | 0.14 | 1.00 | 0.58 | 0.38 | 0.92 |
| pheT | -72828.14 | -72809.01 | 0.02* | 0.03 | 0.15 | 1.00 | 0.59 | 0.37 | 0.92 |
| Orn | -14477.09 | -14474.10 | 0.15 | 0.01 | 0.11 | 1.00 | 0.51 | 0.42 | 0.94 |
| rplR | -6463.44 | -6462.78 | 0.81 | 0.01 | 0.11 | 1.00 | 0.41 | 0.52 | 0.94 |
| gidA | -48523.88 | -48515.69 | 0.02* | 0.01 | 0.10 | 1.00 | 0.57 | 0.39 | 0.94 |
| mraW | -27320.87 | -27316.95 | 0.17 | 0.01 | 0.12 | 1.00 | 0.53 | 0.44 | 0.95 |
| glyS | -58718.79 | -58711.50 | 0.06 | 0.02 | 0.12 | 1.00 | 0.44 | 0.50 | 0.95 |
| rplV | -5196.22 | -5195.46 | 0.82 | 0.02 | 0.19 | 1.00 | 0.82 | 0.17 | 0.95 |
| obgE | -30463.72 | -30460.65 | 0.18 | 0.01 | 0.10 | 1.00 | 0.51 | 0.45 | 0.95 |
| prfA | -28119.78 | -28117.17 | 0.21 | 0.01 | 0.10 | 1.00 | 0.50 | 0.47 | 0.95 |
| aspS | -47605.40 | -47601.81 | 0.32 | 0.02 | 0.13 | 1.00 | 0.49 | 0.43 | 0.95 |
| Frr | -14880.83 | -14879.66 | 0.72 | 0.03 | 0.15 | 1.00 | 0.60 | 0.38 | 0.96 |
| yggX | -6552.94 | -6552.34 | 0.72 | 0.02 | 0.10 | 1.00 | 0.50 | 0.48 | 0.96 |
| engA | -41097.47 | -41093.58 | 0.21 | 0.02 | 0.10 | 1.00 | 0.56 | 0.40 | 0.96 |
| trmE | -38986.82 | -38983.78 | 0.29 | 0.01 | 0.10 | 1.00 | 0.52 | 0.41 | 0.96 |
| dnaN | -30511.33 | -30508.33 | 0.33 | 0.02 | 0.10 | 1.00 | 0.64 | 0.32 | 0.96 |
| Ffh | -33607.23 | -33604.40 | 0.18 | 0.02 | 0.10 | 1.00 | 0.67 | 0.31 | 0.97 |
| rbfA | -11245.17 | -11244.33 | 0.72 | 0.02 | 0.10 | 1.00 | 0.37 | 0.52 | 0.97 |
| Pnp | -49921.49 | -49919.78 | 0.81 | 0.01 | 0.11 | 1.00 | 0.56 | 0.36 | 0.97 |
| infB | -65673.58 | -65671.73 | 0.58 | 0.02 | 0.12 | 1.00 | 0.49 | 0.45 | 0.98 |
| rplJ | -9987.03 | -9986.93 | 0.97 | 0.04 | 0.17 | 1.16 | 0.63 | 0.34 | 0.98 |
| yidC | -43555.73 | -43555.35 | 0.98 | 0.03 | 0.20 | 1.00 | 0.56 | 0.38 | 0.98 |

| | | | | | | | | | |
|---|---|---|---|---|---|---|---|---|---|
| rpoA | -14884.91 | -14884.56 | 0.88 | 0.01 | 0.10 | 1.00 | 0.65 | 0.34 | 1.00 |
| | | | intensified genes | | | | | | |
| Pth | -17673.78 | -17673.35 | 0.83 | 0.01 | 0.10 | 1.00 | 0.43 | 0.53 | 1.00 |
| rplX | -5745.55 | -5745.53 | 0.99 | 0.02 | 0.13 | 1.00 | 0.52 | 0.46 | 1.00 |
| nusA | -34326.92 | -34326.82 | 0.98 | 0.01 | 0.11 | 1.00 | 0.65 | 0.34 | 1.01 |
| ftsH | -42017.26 | -42016.70 | 0.90 | 0.01 | 0.11 | 1.00 | 0.71 | 0.26 | 1.02 |
| yrbA | -7244.76 | -7244.54 | 0.88 | 0.03 | 0.12 | 999.00 | 0.38 | 0.62 | 1.02 |
| rpmA | -4779.06 | -4778.99 | 0.97 | 0.02 | 0.18 | 1.00 | 0.77 | 0.21 | 1.02 |
| rplA | -14992.97 | -14992.61 | 0.88 | 0.02 | 0.13 | 1.00 | 0.60 | 0.38 | 1.02 |
| rpsG | -7517.22 | -7516.85 | 0.90 | 0.01 | 0.10 | 1.00 | 0.71 | 0.26 | 1.03 |
| pyrH | -16848.44 | -16847.17 | 0.58 | 0.01 | 0.10 | 1.00 | 0.68 | 0.31 | 1.03 |
| rpsF | -7439.89 | -7439.63 | 0.93 | 0.04 | 0.22 | 1.00 | 0.49 | 0.43 | 1.04 |
| rpsA | -30783.29 | -30781.23 | 0.96 | 0.02 | 0.14 | 1.00 | 0.64 | 0.35 | 1.04 |
| aceE | -58653.32 | -58648.72 | 0.58 | 0.02 | 0.16 | 1.00 | 0.57 | 0.41 | 1.05 |
| Tsf | -21220.15 | -21218.33 | 0.54 | 0.03 | 0.17 | 1.00 | 0.47 | 0.51 | 1.05 |
| rplQ | -6480.60 | -6479.79 | 0.67 | 0.02 | 0.10 | 1.00 | 0.51 | 0.39 | 1.05 |
| rplO | -8749.55 | -8748.57 | 0.82 | 0.01 | 0.12 | 1.00 | 0.43 | 0.54 | 1.06 |
| rplT | -6179.76 | -6178.22 | 0.44 | 0.01 | 0.10 | 1.00 | 0.62 | 0.36 | 1.06 |
| secY | -25094.28 | -25083.52 | 0.02* | 0.01 | 0.10 | 1.00 | 0.64 | 0.36 | 1.06 |
| rplK | -7959.80 | -7958.73 | 0.72 | 0.02 | 0.15 | 1.00 | 0.68 | 0.31 | 1.06 |
| rplI | -10623.93 | -10622.72 | 0.64 | 0.03 | 0.15 | 1.00 | 0.39 | 0.57 | 1.06 |
| rplE | -8438.27 | -8437.15 | 0.72 | 0.02 | 0.17 | 1.00 | 0.64 | 0.34 | 1.07 |
| rpsR | -3131.34 | -3130.83 | 0.82 | 0.01 | 0.13 | 1.00 | 0.64 | 0.33 | 1.08 |
| rho | -21447.11 | -21440.78 | 0.02* | 0.01 | 0.10 | 1.00 | 0.84 | 0.16 | 1.08 |
| rplM | -7428.74 | -7427.16 | 0.44 | 0.02 | 0.17 | 1.00 | 0.62 | 0.35 | 1.09 |
| rpsB | -13719.86 | -13716.33 | 0.27 | 0.02 | 0.14 | 1.00 | 0.55 | 0.39 | 1.09 |
| rpsP | -5636.77 | -5635.06 | 0.44 | 0.02 | 0.13 | 1.00 | 0.43 | 0.48 | 1.10 |
| rplP | -6088.36 | -6085.79 | 0.22 | 0.02 | 0.14 | 2.79 | 0.65 | 0.35 | 1.11 |
| rplS | -6084.20 | -6079.72 | 0.18 | 0.02 | 0.10 | 1.00 | 0.53 | 0.43 | 1.11 |
| rplB | -13727.93 | -13721.89 | 0.06 | 0.01 | 0.14 | 1.00 | 0.60 | 0.37 | 1.13 |
| rpsM | -6351.64 | -6349.60 | 0.33 | 0.02 | 0.23 | 1.00 | 0.57 | 0.38 | 1.13 |
| infA | -3065.84 | -3062.17 | 0.08 | 0.01 | 0.10 | 28.43 | 0.71 | 0.28 | 1.14 |
| rpoB | -73750.02 | -73701.12 | 0.02* | 0.01 | 0.17 | 1.00 | 0.66 | 0.32 | 1.16 |
| rplN | -4879.27 | -4875.46 | 0.06 | 0.01 | 0.13 | 1.00 | 0.64 | 0.33 | 1.21 |
| rpoC | -78720.19 | -78645.42 | 0.02* | 0.01 | 0.19 | 1.00 | 0.69 | 0.28 | 1.21 |
| rpsN | -5546.14 | -5539.38 | 0.04* | 0.02 | 0.17 | 1.00 | 0.51 | 0.47 | 1.23 |
| rpsJ | -3591.13 | -3578.45 | 0.02* | 0.01 | 0.10 | 1.00 | 0.78 | 0.21 | 1.36 |
